# LncRNA BRE-AS1 Regulates the JAK2/STAT3-mediated Inflammatory Activation via the miR-30b-5p/SOC3 Axis in THP-1 cells

**DOI:** 10.1101/2024.03.06.583653

**Authors:** Jae-Joon Shin, Kyoungho Suk, Won-Ha Lee

## Abstract

Long non-coding RNAs (lncRNAs) have emerged as pivotal regulators in numerous biological processes, including macrophage-mediated inflammatory responses, which play a critical role in the progress of diverse diseases. This study focuses on the regulatory function of lncRNA BRE-AS1 in modulating the inflammatory activation of monocytes/macrophages. Employing the THP-1 cell line as a model, we demonstrate that lipopolysaccharide (LPS) treatment significantly upregulates BRE-AS1 expression. Notably, specific knockdown of BRE-AS1 via siRNA transfection enhances LPS-induced expression of interleukin (IL)-6 and IL-1β, while not affecting tumor necrosis factor (TNF)-α levels. This selective augmentation of pro-inflammatory cytokine production coincides with increased phosphorylation of JAK2 and STAT3. Furthermore, BRE-AS1 suppression results in the downregulation of SOCS3, an established inhibitor of the JAK2/STAT3 pathway. Bioinformatics analysis identified binding sites for miR-30b-5p on both BRE-AS1 and SOCS3 mRNA. Intervention with a miR-30b-5p inhibitor and a synthetic RNA fragment that represents the miR-30b-5p binding site on BRE-AS1 attenuates the pro-inflammatory effects of BRE-AS1 knockdown. Conversely, a miR-30b-5p mimic replicated the BRE-AS1 attenuation outcomes. Our findings elucidate the role of lncRNA BRE-AS1 in modulating inflammatory activation in THP-1 cells via the miR-30b-5p/SOCS3/JAK2/STAT3 signaling pathway, proposing that manipulation of macrophage BRE-AS1 activity may offer a novel therapeutic avenue in diseases characterized by macrophage-driven pathogenesis.

## Introduction

The Janus kinase 2 (JAK2) plays a multifaceted and pivotal role in various physiological processes, including immune regulation, cell differentiation, and apoptosis [1, 2]. As a substrate of JAK2, the signal transducer and activator of transcription 3 (STAT3) functions as an intracellular cytoplasmic transcription factor, actively participating in a wide range of biological processes including cell growth, survival, differentiation, and immune responses [3, 4]. The JAK2/STAT3 signaling pathway is crucial for lipopolysaccharide (LPS)-induced inflammation, where LPS-induced Toll-like receptor 4 (TLR4) signaling initiates JAK2 phosphorylation, subsequently leading to the phosphorylation of STAT3 on its tyrosine residues [5]. Recent studies have demonstrated the essential roles of JAK2 and STAT3 in regulating the expression of IL-6, and IL-1β in macrophages stimulated by LPS [6-8]. The suppressor of Cytokine Signaling 3 (SOCS3), a member of the SOCS family, plays a significant role in regulating cell signal transduction pathways, being inducible by various cytokines and pro-inflammatory factors [9]. SOCS3 suppresses the JAK/STAT pathway by interacting with both the JAK kinase and cytokine receptors, thus preventing STAT3 phosphorylation [10, 11]. Additionally, LPS triggers SOCS3 expression via multiple signaling pathways, with SOCS3 acting as a negative feedback regulator in macrophages, especially in the IL-6 and gp130-mediated pathways [12]. Recent advances in RNA biology have further shown that miRNA, such as miR-30b-5p, can regulate SOCS3 expression in LPS-stimulated macrophages [13, 14].

Long non-coding RNAs (lncRNAs), characterized as transcripts longer than 200 base pairs and largely lacking protein-coding capability, play crucial roles in various biological processes through interactions with RNA, DNA, and proteins [15, 16]. These interactions enable lncRNAs to regulate transcriptional, post-transcriptional, translational, and epigenetic modification processes under diverse conditions, thereby contributing to cellular function and homeostasis [17, 18]. Recent research has highlighted the significance of lncRNAs in modulating immune functions, including inflammation [19]. One such lncRNA, BRE-AS1 (also known as BABAM2-AS1), spans 1.6 kilobases and is located on chromosome 2p23.2. It has garnered attention in the study of various diseases for its ability to influence cellular responses through interactions with miRNAs and proteins. Specifically, BRE-AS1 has been shown to regulate prostate cancer (PC) cell proliferation and apoptosis by upregulating miR-145-5p, while in triple-negative breast cancer (TNBC), it hinders cell proliferation, migration, and invasion through the downregulation of miR-21 [20, 21]. Furthermore, BRE-AS1 impedes the growth and survival of non-small cell lung cancer (NSCLC) cells through STAT3 [22]. Additionally, high expression levels of BRE-AS1 have been observed in bone marrow, suggesting its potential involvement in hematopoiesis or immune regulation [13]. However, research into the precise function of BRE-AS1 in regulating macrophage inflammatory responses remains unexplored.

In our study, we sought to investigate the role of BRE-AS1 in TLR4-induced inflammatory activation within the human monocytic leukemia cell line THP-1. Drawing on bioinformatic analyses and prior research, we hypothesized that BRE-AS1 influences the inflammatory response by mediating the regulation of SOCS3 via the sequestration of miR-30b-5p. To probe this hypothesis, we executed a series of experiments using synthetic RNA molecules. These experiments were designed to elucidate the mechanistic pathways through which BRE-AS1 potentially alters the inflammatory landscape, particularly focusing on its interaction with miR-30b-5p and the subsequent impact on SOCS3 expression. Through these investigations, we aimed to provide new insights into the regulatory functions of lncRNAs in inflammation and highlight the therapeutic potential of targeting BRE-AS1 in inflammatory diseases.

## Materials and methods

### Cell culture and reagents

The THP-1 cell line was cultured in RPMI 1640 medium (WelGENE Inc., Daegu, Korea), enriched with 10% fetal bovine serum (FBS), 0.05□mM β-mercaptoethanol, glucose, and streptomycin-penicillin, maintained at 37□°C in a 5% CO2 atmosphere. Rabbit monoclonal antibodies (mAbs) targeting STAT3, phospho-STAT3 (Tyr705), and JAK2 were sourced from Cell Signaling Technology (Danvers, MA). Mouse mAb against β-actin was procured from Santa Cruz Biotechnology (Dallas, TX). Mouse mAbs for SOCS3 were acquired from both Santa Cruz Biotechnology (Dallas, TX) and Abcam (Cambridge, U.K.). Rabbit mAb directed at phospho-JAK2 (Tyr1007/1008) was obtained from Invitrogen (Eugene, OR). Bacterial LPS and N-acetylcysteine (NAC) were purchased from Sigma-Aldrich (St. Louis, MO). The DharmaFECT 1 small interfering RNA (siRNA) transfection reagent was acquired from Dharmacon (Lafayette, CO). Scramble siRNA, siRNAs targeting BRE-AS1, fragments of BRE-AS1, and the miR-30b-5p inhibitor were supplied by Bioneer (Daejeon, Korea).

### Cell transfection

THP-1 cells (3.0□×□10^5^ cells) were initially seeded in 6-well plates using an antibiotic-free culture medium. After 18 hours, the cells underwent transfection with siRNA at a concentration of 100 nM, alongside decoy RNA, microRNA inhibitor, and microRNA mimic, each at a concentration of 200 nM. This transfection used DharmaFECT 1 siRNA transfection reagent, adhering to the manufacturer’s guidelines. The transfected cells were then harvested for mRNA and protein level analysis 24 hours post-transfection. The sequences for BRE-AS1 siRNA were as follows: siRNA (sense, 5’-GUUGUUGUGAGGACUAAAUGA-3’ and antisense, 5’-AUUUAGUCCUCACAACAACCC-3’) and decoy RNA (5’-CGGGGUUUACAGGAA-3’).

### Quantitative real-time PCR (qRT-PCR)

Total cellular RNA was extracted employing TRIzol Reagent. Subsequently, the isolated RNAs underwent treatment with RNase-free DNase I (Takara Bio, Otsu, Shiga, Japan) to remove any contaminating DNA. Following this treatment, the RNAs were utilized for cDNA synthesis, which was performed using the Reverse Transcription Master Premix (Elpis Biotech, Daejeon, Korea). Quantitative RT-PCR (qRT-PCR) analyses were conducted on a StepOnePlus system (Applied Biosystems, Foster City, CA) employing SYBR Premix Ex Taq (Takara Bio), and the specific primer sequences used are detailed in Table I. The threshold cycle (Ct) values obtained from each reaction were normalized against the GAPDH Ct values to account for variations in sample loading and to ensure accurate quantification.

**Table □.**
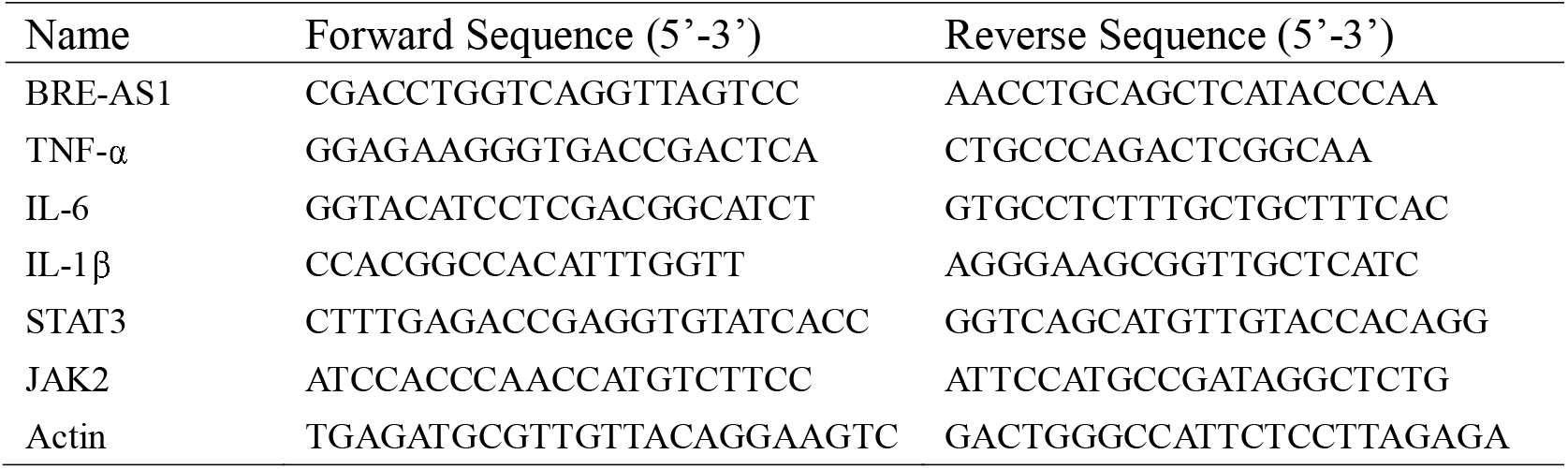
Primers used for quantitative reverse-transcription PCR.

### Enzyme-linked immunosorbent assay (ELISA)

The concentrations of cytokines in culture supernatants were quantified using ELISA Kits (Invitrogen, Biolegend). The ELISA procedure was carried out strictly following the manufacturer’s instructions. Colorimetric changes indicative of cytokine concentrations were detected using a microplate reader, calibrated to a wavelength of 450 nm with a correction for absorption at 540 nm. To ensure accuracy and reproducibility, measurements were conducted in triplicate.

### Western blot

Cell pellets were lysed using NP-40 (IGEPAL CA-630) lysis buffer (150 mM NaCl, 1% IGEPAL CA-630, 50 mM Tris, pH 8.0), supplemented with a protease inhibitor cocktail (Calbiochem, San Diego, CA) and a phosphatase inhibitor cocktail (Sigma-Aldrich). The lysate was clarified by centrifugation at 12,000 rpm for 15 minutes at 4°C, removing cellular debris. The supernatant, containing the proteins, was treated with 100 mM dithiothreitol (DTT) and heated to denature the proteins. Subsequently, protein samples were subjected to electrophoresis on a 10% or 12% polyacrylamide gel and transferred onto a PVDF membrane (Milipore, Burlington, USA). The membrane was then blocked with a 5% bovine serum albumin (BSA) solution in TBS containing 0.1% Tween 20 (TBST) for 1 hour. Following three washes with TBST, the membrane was incubated overnight at 4°C with primary antibodies diluted in the blocking solution. After additional washes with TBST, the membrane was incubated with horseradish peroxidase (HRP)-conjugated secondary antibodies at 4°C for 1 hour. Following further TBST washes, chemiluminescent detection was performed using a detection reagent (Corebio, Seoul, Korea).

### Bioinformatic analysis

the potential interaction site between BRE-AS1 and miR-30b-5p was investigated using DIANA tools, accessible at https://diana.e-ce.uth.gr/lncbasev3/interactions. Furthermore, the binding site prediction for the interaction between miR-30b-5p and SOCS3 was conducted using the miRDB online database, available at https://mirdb.org/mirdb.

### Statistical analysis

Statistical analyses were conducted using two-way ANOVA for comparisons among multiple groups, with Bonferroni post-tests applied for detailed comparisons between specific groups. For direct comparisons between two distinct groups, an independent samples unpaired Student’s t-test was employed. A threshold of P<0.05 was set to denote statistical significance. All experimental procedures were performed in triplicate and repeated more than three times to ensure reliability. The results are presented as the mean ± standard error of the mean (SEM).

## Results

### The reduction of BRE-AS1 expression results in increased IL-6 and IL-1β expression in activated THP-1 cells

To elucidate the role of BRE-AS1 in monocyte/macrophage inflammation, we initially examined alterations in BRE-AS1 expression levels in THP-1 cells following LPS stimulation. The levels of BRE-AS1, alongside TNF-α mRNA as a positive control, peaked at the 2-hour mark post-stimulation, subsequently showing a gradual decline over time (Fig 1A and B). This observation suggests a dynamic response of BRE-AS1 to inflammatory stimuli, indicating its potential involvement in the early phase of the inflammatory response in monocytes/macrophages.

**Figure 1.**
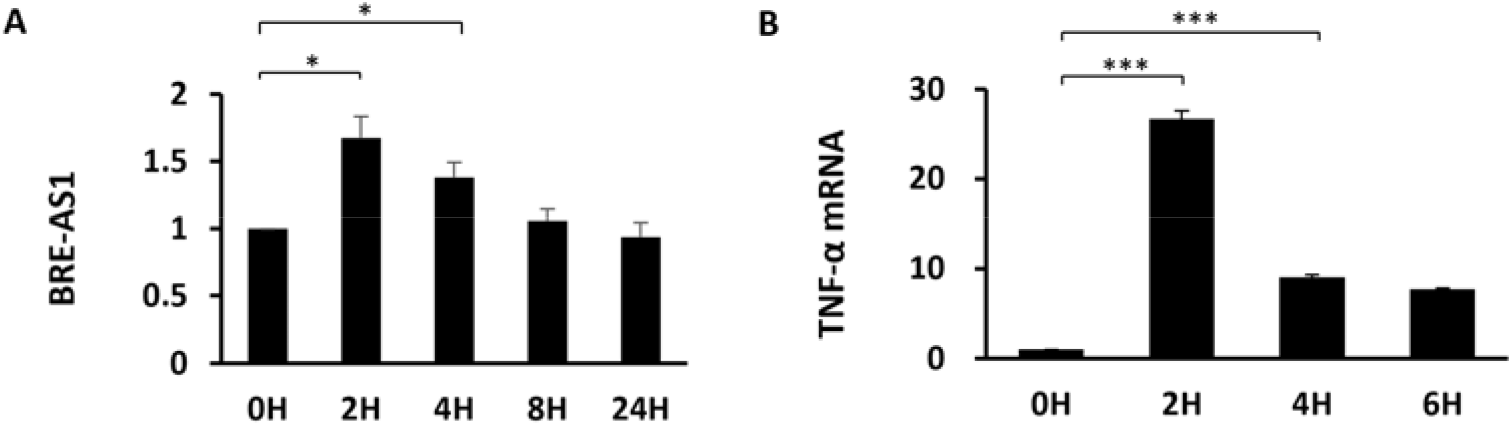
Upregulation of BRE-AS1 Expression Following LPS Treatment. (A, B) THP-1 cells were treated with LPS (1000 ng/ml) for specified durations. The expression levels of BRE-AS1 (A) and TNF-α mRNA (B) were quantified using real-time PCR. *p<0.05; ***p<0.001.

To explore the role of BRE-AS1 in LPS-triggered inflammatory responses, we employed specific siRNA to reduce BRE-AS1 expression (Fig 2A). Subsequent LPS stimulation of THP-1 cells revealed that diminishing BRE-AS1 expression led to increased mRNA levels of IL-6 and IL-1β, whereas TNF-α levels were unaffected (Fig 2B-D). Consistent with these findings, the attenuation of BRE-AS1 amplified the LPS-induced secretion of IL-6, while the secretion levels of TNF-α remained stable (Fig 2E and F). These results underline the specific regulatory role of BRE-AS1 in modulating the inflammatory response, particularly influencing the expression and secretion of key cytokines such as IL-6 and IL-1β in the context of LPS stimulation.

**Figure 2.**
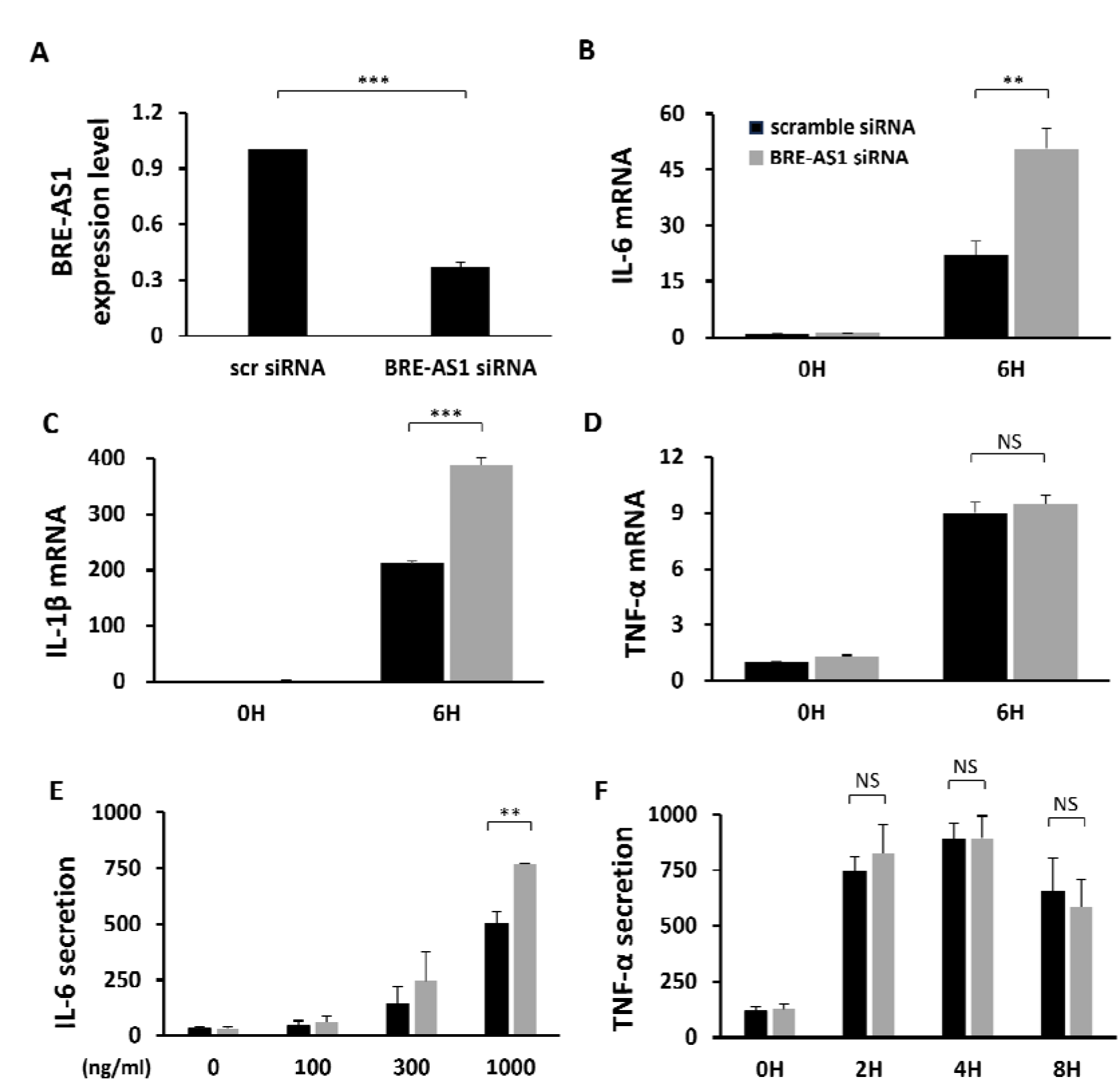
Reduction of BRE-AS1 Enhances IL-6 and IL-1β Expression Without Affecting TNF-α. (A) The effectiveness of the knockdown was evaluated by real-time PCR 24 hours after transfecting THP-1 cells with scramble or BRE-AS1 siRNA. (B-D) Following LPS stimulation (1000 ng/ml), mRNA levels of cytokines were quantified using real-time PCR. (E, F) Cytokine secretion levels were measured by ELISA after LPS stimulation for 24 hours at specified concentrations for IL-6 (E) and at indicated durations for TNF-α with LPS (1000 ng/ml) (F). **p<0.01; ***p<0.001.

### BRE-AS1 enhances the expression of pro-inflammatory cytokines through the SOCS3/JAK2/STAT3 pathway

STAT3 is a critical regulator that enhances IL-1β and IL-6 expression in macrophages stimulated by LPS, independent of TNF-α expression [6-8]. Stimulation by LPS leads to the phosphorylation of JAK2, which subsequently activates STAT3 [4]. SOCS3 acts as an upstream regulator that inhibits the JAK2/STAT3 pathway and is upregulated in response to LPS stimulation in macrophages [12, 23]. Drawing on the findings presented in Figure 2, we proposed that BRE-AS1 might target the STAT3 pathway and its upstream regulators. To explore this hypothesis, we reduced BRE-AS1 expression and then evaluated the expression levels and/or activation status of JAK2, STAT3, and SOCS3.

THP-1 cells were transfected with BRE-AS1 siRNA and subsequently stimulated with LPS. The protein levels and phosphorylation status of signaling molecules were assessed using Western blot analysis. Following the attenuation of BRE-AS1, a decrease in the SOCS3 protein level was observed. However, the phosphorylation levels of JAK2 and STAT3 were increased without significant alterations in the total protein levels (Fig 3A and B). Correspondingly, the mRNA level of SOCS3 decreased (Fig 3C), while there were no significant changes in the mRNA levels of JAK2 and STAT3 (Fig 3D and E). This suggests that the modulation of BRE-AS1 directly impacts the SOCS3 protein expression and the phosphorylation state of JAK2 and STAT3.

**Figure 3.**
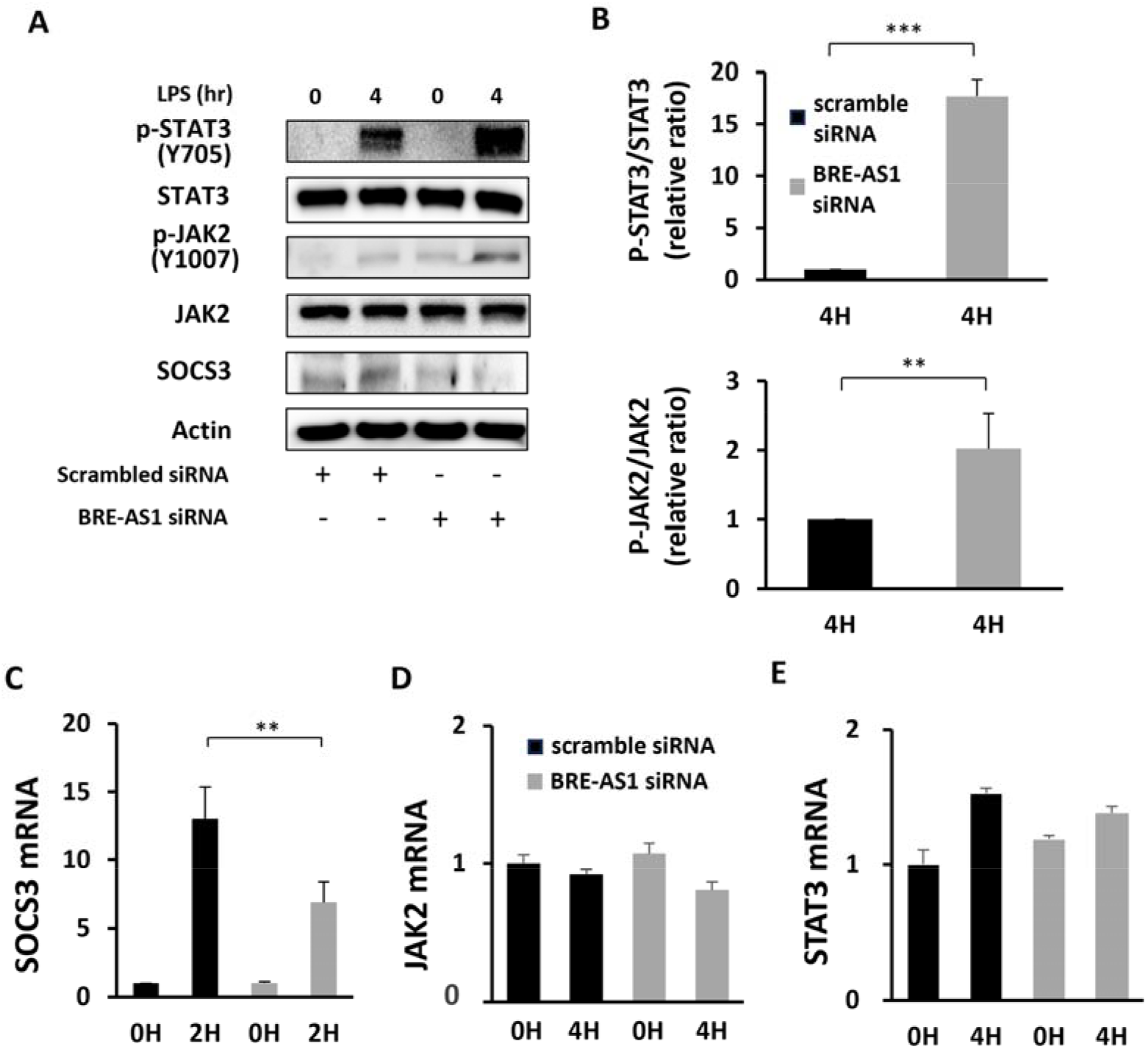
Inhibition of BRE-AS1 Enhances STAT3 Phosphorylation and IL-6 Expression via the SOCS3/JAK2/STAT3 Pathway. (A-E) THP-1 cells, after being transfected with BRE-AS1 siRNA, were stimulated with LPS (1000 ng/ml). Protein levels of SOCS3, JAK2, p-JAK2, STAT3, and p-STAT3 were analyzed by Western blot in cell extracts. (B) Protein expression levels in (A) were quantified using ImageJ software. (C-E) mRNA levels of SOCS3, JAK2, and STAT3 were measured using real-time PCR. **p<0.01; ***p<0.001.

### BRE-AS1 regulates SOCS3 mRNA levels through miR-30b-5p in THP-1 cells

The mechanism of lncRNAs modulating gene expression often involves their competitive binding to miRNAs, thereby inhibiting miRNA function [24]. The DIANA Tools database (https://diana.e-ce.uth.gr/lncbasev3/interactions) indicated a significant potential for interaction between BRE-AS1 and miR-30b-5p. MiR-30b-5p is known to target SOCS3 mRNA, affecting its expression in conditions such as acute lung injury and in LPS-stimulated Raw264.7 cells [13, 14]. Based on this, it is hypothesized that BRE-AS1 may target miR-30b-5p, subsequently modulating the SOCS3/JAK2/STAT3 signaling pathway.

The potential interaction sites among BRE-AS1, miR-30b-5p, and SOCS3 were predicted using DIANA tools [13] and miRDB [25] (Fig 4A). To investigate if BRE-AS1 modulates its effects through the sequestration of miR-30b-5p, a decoy RNA mimicking the miR-30b-5p binding site on BRE-AS1 was utilized. Decoy RNA, comprising artificially synthesized oligonucleotide fragments, possesses a complementary site to the target miRNA, thereby inhibiting its function [26]. The reduction of BRE-AS1 via siRNA transfection diminishes its capacity to act as a sponge for miR-30b-5p, potentially increasing miR-30b-5p levels. Consequently, transfecting cells with decoy RNA aims to offset the decreased presence of BRE-AS1. As anticipated, the alterations in SOCS3, IL-6, and IL-1β mRNA levels prompted by the reduction of BRE-AS1 were mitigated by the decoy RNA (Fig 4B-D). This compensatory effect was similarly observed in the secretion of IL-6 (Fig 4E). Additionally, the protein-level modifications in JAK2 and STAT3 induced by BRE-AS1 siRNA transfection were reversed upon introducing the decoy RNA (Fig 4F), underscoring the functional importance of BRE-AS1 in regulating these molecular interactions through miR-30b-5p.

**Figure 4.**
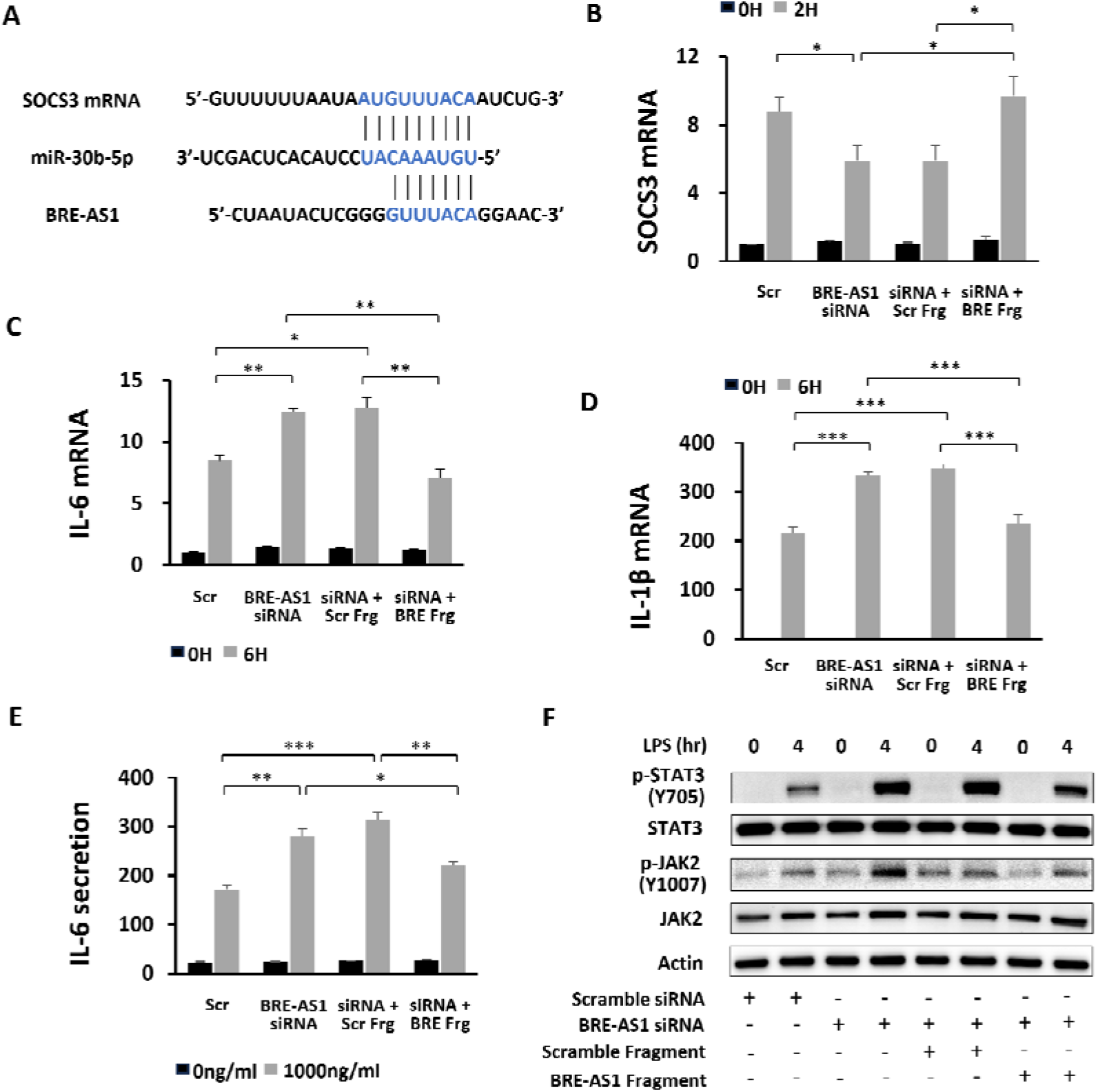
BRE-AS1 Modulation of SOCS3 mRNA via miR-30b-5p Interaction. (A) Identification of complementary binding sites between BRE-AS1, miR-30b-5p, and SOCS3 was performed. (B-F) THP-1 cells were transfected with BRE-AS1 siRNA and a decoy RNA (BRE-AS1 fragment) for 24 hours before being stimulated with LPS (1000 ng/ml). (B-D) mRNA expression levels of SOCS3, IL-6, and IL-1β were quantified using real-time PCR. (E) IL-6 secretion levels were determined by ELISA. (F) Protein expression in the JAK2/STAT3 pathway was analyzed by Western blot. *p<0.05; **p<0.01; ***p<0.001.

To further validate the BRE-AS1/miR-30b-5p/SOCS3 axis, additional experiments were conducted in THP-1 cells using a miR-30b-5p inhibitor. The application of the miR-30b-5p inhibitor mitigated the effects of siRNA on the mRNA levels of SOCS3 and IL-6 (Fig 5A and B). It also counteracted protein-level changes within the SOCS3/JAK2/STAT3 pathway caused by siRNA (Fig 5C). Moreover, to ascertain miR-30b-5p’s role in regulating SOCS3 expression in THP-1 cells, a miR-30b-5p mimic was employed. Introducing the miR-30b-5p mimic led to decreased mRNA level of SOCS3 while those of IL-6, and IL-1β were increased (Fig 5D-F). Accordingly, SOCS3 protein level was decreased while phosphorylation levels of JAK2 and STAT3 were increased (Fig 5G). These findings collectively underscore that BRE-AS1 modulates the SOCS3/JAK2/STAT3 signaling pathway via miR-30b-5p, subsequently affecting the expression of IL-6 and IL-1β. This elucidates the intricate regulatory mechanisms through which BRE-AS1 influences inflammatory responses, highlighting its potential as a therapeutic target in inflammation-related diseases.

**Figure 5.**
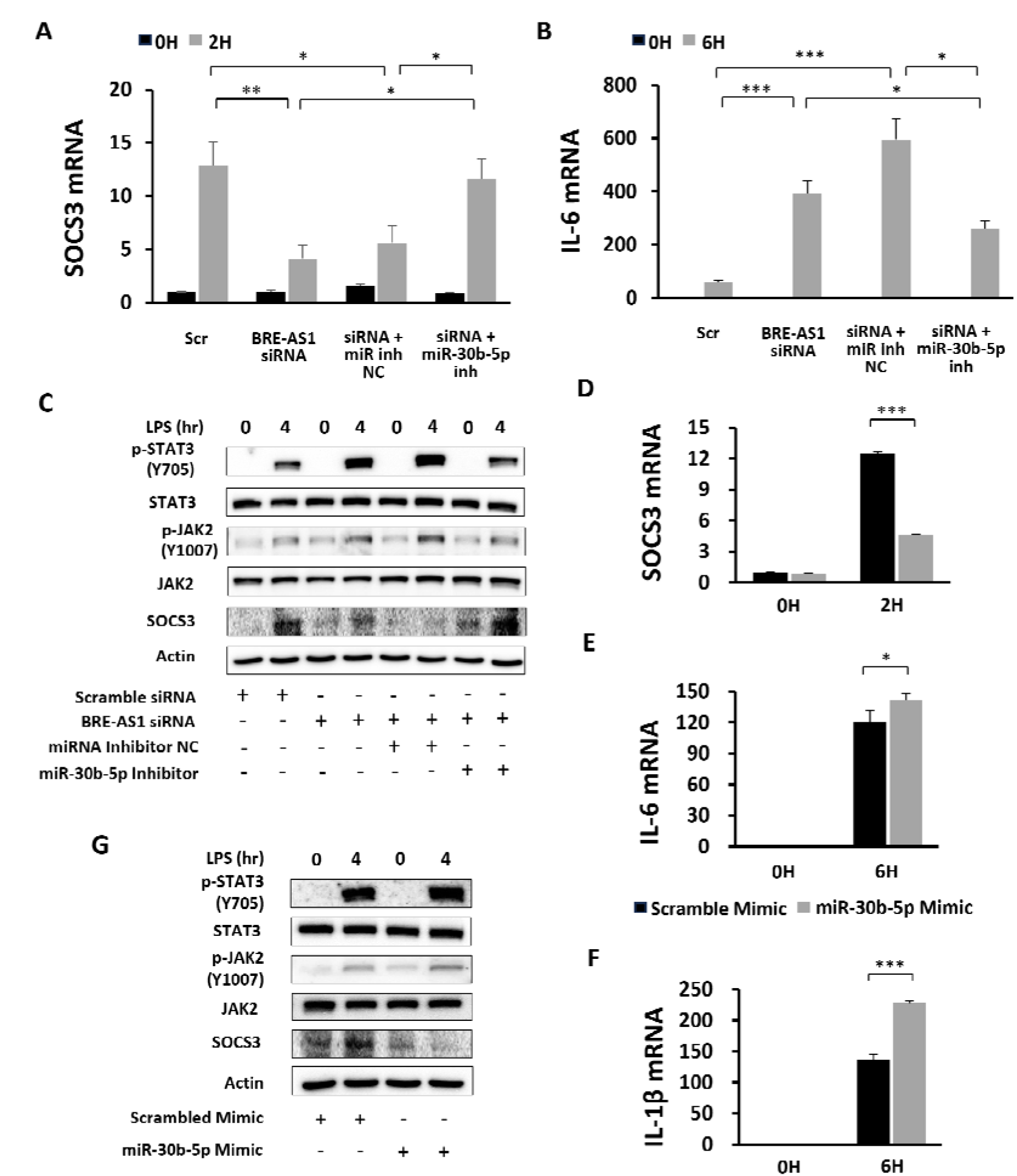
BRE-AS1 Modulation of SOCS3 mRNA via miR-30b-5p Interaction. (A-C) THP-1 cells were transfected with a miR-30b-5p inhibitor alongside BRE-AS1 siRNA for 24 hours before LPS stimulation (1000 ng/ml). (A, B) mRNA levels of SOCS3 and IL-6 were quantified using real-time PCR. (C) Protein levels within the SOCS3/JAK2/STAT3 pathway were analyzed by Western blot. (D-G) THP-1 cells underwent transfection with miR-30b-5p mimic and mimic control for 24 hours. (D-F) The mRNA expression of SOCS3, IL-6, and IL-1β was assessed using real-time PCR. (G) Protein levels of the SOCS3/JAK2/STST3 pathway were evaluated by Western blot. *p<0.05; **p<0.01; ***p<0.001.

## Discussion

Inflammation is a critical component of the innate immune defense mechanism against harmful stimuli [27]. However, dysregulated or inappropriate inflammatory responses are implicated in the pathogenesis of various diseases, such as atherosclerosis, systemic lupus erythematosus, and rheumatoid arthritis [7, 28]. This underscores the importance of research in this area. Within the realm of inflammation, recent studies have explored the role of lncRNAs in regulating biological pathways across multiple diseases [29]. Specifically, BRE-AS1, known for its high expression levels in bone marrow, interacts with proteins or miRNAs that regulate inflammatory signaling pathways [13, 22]. This led us to explore the potential impact of BRE-AS1 on macrophages and to investigate its role in macrophage inflammatory activation, along with the underlying mechanisms. Our current findings introduce a novel aspect of gene regulation, indicating that BRE-AS1 may affect the inflammatory response through the miR-30b-5p-mediated regulation of SOCS3 expression, which in turn influences the JAK2/STSAT3 signaling pathway leading to the production of pro-inflammatory cytokines such as IL-6 and IL-1β (Fig 6). turn affects the levels of IL-6 and IL-1β expression.

**Figure 6.**
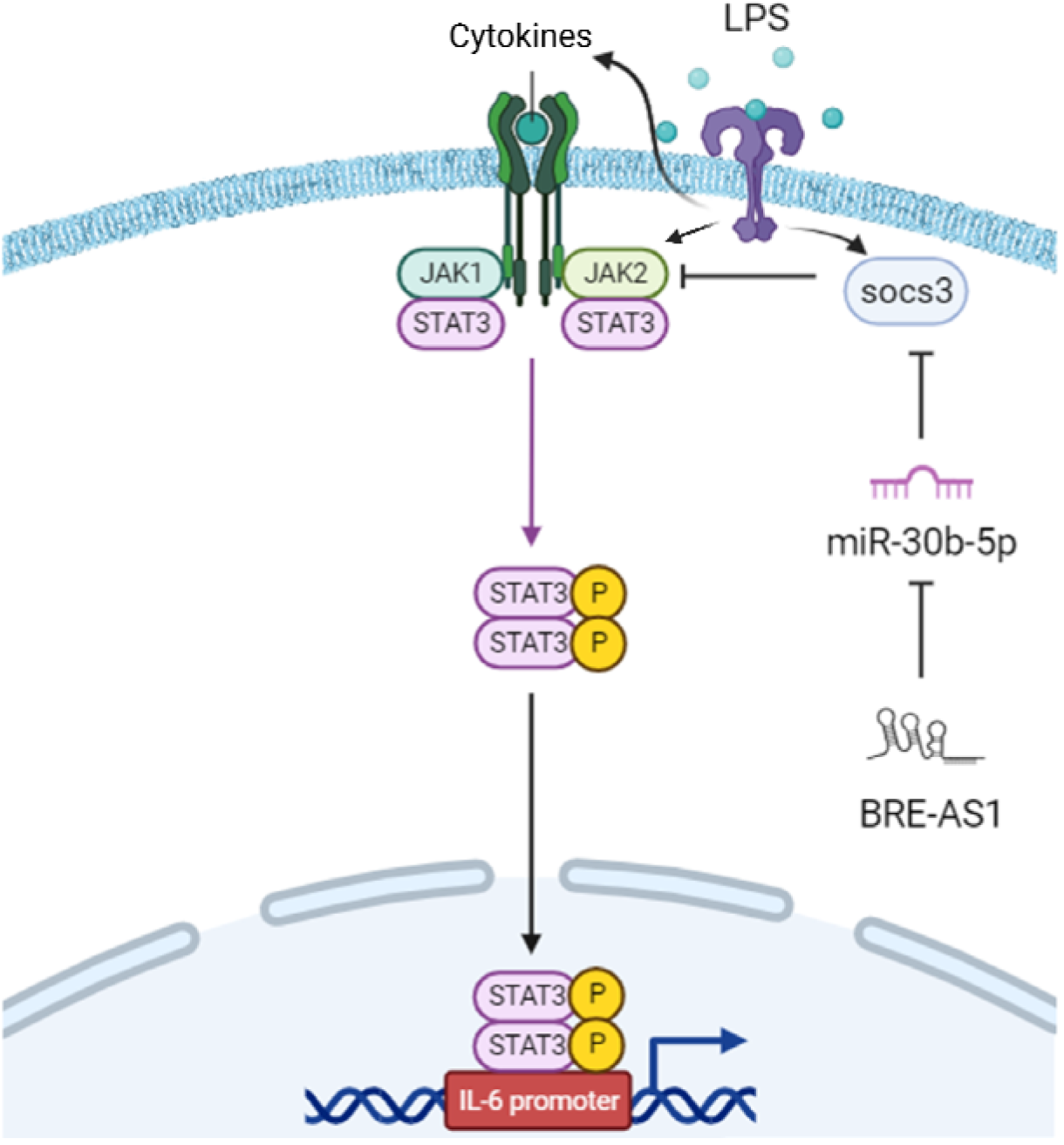
BRE-AS1 regulates macrophage inflammatory activation via the miR-30b-5p/SOCS3/JAK2/STAT3 Pathway. LncRNA BRE-AS1 modulates SOCS3 expression by serving as a competitive endogenous RNA for miR-30b-5p. SOCS3 subsequently controls the phosphorylation and activation of JAK2 and STAT3, which in turn affects the levels of IL-6 and IL-1 β expression.

Knocking down BRE-AS1 led to an increase in the expression of IL-6 and IL-1β following LPS stimulation in THP-1 cells, while TNF-α expression remained unchanged. The transcription factor NF-κB, activated by signals from TLR4, is recognized as a crucial regulator of inflammatory cytokine production and inflammatory activity. NF-κB controls the expression of pro-inflammatory cytokines, including TNF-α, IL-1, and IL-6, in macrophages [30]. However, our findings indicate an elevation in IL-6 and IL-1β expression without significant changes in TNF-α levels after BRE-AS1 knockdown, suggesting that BRE-AS1 may modulate macrophage inflammatory activity through mechanisms that do not solely rely on the regulatory pathways associated with the transcription factor NF-κB following LPS-TLR4 signaling activation. Recent studies have highlighted that the phosphorylation of JAK2 and STAT3 plays a critical role in the expression of IL-6 and IL-1β, independently of TNF-α, in macrophages following LPS stimulation [6-8]. LPS-triggered TLR4 signaling leads to JAK2 phosphorylation, which in turn induces STAT3 phosphorylation. Activated STAT3 then translocates to the nucleus and interacts with specific target genes, modulating inflammatory pathways. We hypothesized that BRE-AS1 might regulate the JAK2/STAT3 pathway, potentially influencing cytokine production. Following BRE-AS1 knockdown, we observed no significant changes in JAK2 and STAT3 expression levels but a significant increase in their phosphorylation levels. These results suggest that BRE-AS1 could impact the activity of JAK2/STAT3 independently of their expression, highlighting a potential mechanism through which BRE-AS1 influences macrophage inflammatory responses.

SOCS3 is crucial in regulating cellular signaling pathways, especially by inhibiting the JAK/STAT pathway [11, 23]. It interacts with JAK kinase and cytokine receptors, thereby preventing STAT3 phosphorylation. MiR-30b-5p’s regulatory role in controlling SOCS3 expression, particularly during acute lung injury, has been studied [14]. Overexpression of miR-30b-5p in RAW264.7 cells leads to decreased SOCS3 mRNA expression, demonstrating a direct regulatory interaction between miR-30b-5p and SOCS3 [14]. Additionally, database analysis revealed a significant potential for interaction between BRE-AS1 and miR-30b-5p. Based on these findings, it was hypothesized that BRE-AS1 modulates SOCS3 through miR-30b-5p, affecting the JAK2/STAT3 pathway’s activation. Upon LPS stimulation in THP-1 cells, BRE-AS1 expression increases, possibly leading to enhanced sequestration of miR-30b-5p and, consequently, liberating SOCS3 mRNA from the miRNA’s inhibitory effects. Therefore, the knockdown of BRE-AS1 is anticipated to raise miR-30b-5p levels, resulting in a decrease in SOCS3 levels. Indeed, the knockdown of BRE-AS1 led to a decrease in SOCS3 expression, confirming SOCS3 as a target of BRE-AS1 regulation. Moreover, the application of synthesized RNAs, including a decoy RNA representing the miR-30b-5p binding site of BRE-AS1 and miR-30b-5p inhibitor/mimic, further confirmed that BRE-AS1 regulates SOCS3 via miR-30b-5p.

Our findings underscore the potential therapeutic value of targeting the BRE-AS1/miR-30b-5p/SOCS3 axis in the treatment of inflammatory conditions. By manipulating this regulatory pathway, it may be possible to develop novel strategies for the management of inflammatory diseases, offering more precise and targeted approaches compared to current therapies.

While this study provides significant insights into the regulation of inflammation via lncRNAs, several limitations must be acknowledged. The use of THP-1 cells as a model system, though informative, necessitates validation of these findings in primary human macrophages and in vivo models to better understand the clinical relevance of this regulatory axis.

Future research should aim to explore the broader implications of BRE-AS1 regulation in different inflammatory contexts and disease models. Investigating the interaction of BRE-AS1 with other components of the inflammatory response and its role in other cell types involved in inflammation could provide a more comprehensive understanding of its function in immune regulation.

## Conclusion

In conclusion, our research identifies a novel regulatory axis involving BRE-AS1, miR-30b-5p, and SOCS3 that modulates inflammatory responses in THP-1 cells. By influencing the JAK2/STAT3 pathway via SOCS3 expression regulation, BRE-AS1 governs the inflammatory process, notably affecting the production of the key pro-inflammatory cytokines IL-6 and IL-1β. This investigation enhances our comprehension of the molecular mechanisms that control inflammatory processes and underscores the therapeutic potential of targeting lncRNAs in treating inflammatory diseases where macrophages play a pivotal role. Further studies are necessary to clarify the clinical implications of this regulatory axis and to investigate its therapeutic applicability in managing inflammatory disorders.

## Declarations

During the preparation of this work the authors used ChatGPT to improve the quality of English writing. After using this service, the authors reviewed and edited the content as needed and takes full responsibility for the content of the publication.

## Author contributions

JJS: Conceptualization, Investigation, Writing-original draft & editing. KS: Conceptualization, Writing-review & editing. WHL: Supervision, validation, Writing-review & editing. All authors read and approved the submitted version.

## Ethical approval

Not applicable

## Consent to participate

Not applicable.

## Consent to publication

Not applicable.

## Availability of data and materials

Not applicable.

## Abbreviations

LncRNA: long noncoding RNA
miRNA: microRNA
JAK2: janus kinase 2
p-JAK2: phospho-JAK2
STAT3: Signal transducer and activator of transcription 3
p-STAT3: phospho-STAT3
SOCS3: Suppressor Of Cytokine Signaling 3
LPS: lipopolysaccharide
TLR4: toll-like receptor 4
IL-6: interleukin-6
IL-1β: interleukin 1 beta
TNF-α: tumor necrosis factor-α
siRNA: small interfering RNA
PC: prostate cancer
TNBC: triple-negative breast cancer
NSCLC: non-small-cell lung cancer
FBS: fetal Bovine Serum

